# A CD25-CCR7 complex initiates non-canonical IL-2 signaling

**DOI:** 10.1101/2025.02.03.636356

**Authors:** Sarah Hyun Ji Kim, Hosup Lee, Alexandre Gingras, Klaus Ley, Jamie B. Spangler, Mark H. Ginsberg

## Abstract

IL-2, a central regulator of immune function, binds to its receptor subunit CD25 (IL-2Rα), promoting IL-2 interaction with β and γ subunits to trigger the canonical IL-2 signaling pathway. An anti-mouse CD25 antibody, PC61, triggers alternative IL-2 signaling, leading to integrin activation. PC61 induces a complex formed by the IL-2-dependent association of CD25 with CCR7, suggesting that the formation of this complex initiates alternative IL-2 signaling. Here, we used structure-based design together with combinatorial screening to identify an IL-2 mutant (denoted IL-2(E52K)) that spares canonical IL-2 signaling but disrupts both PC61-induced complex formation and integrin activation while retaining the full CD25 affinity of the parent molecule. We also report that heparan sulfate (HS), a physiological ligand of IL-2 that triggers alternative signaling, induced IL-2-dependent CD25-CCR7 association, whereas IL-2(E52K) failed to support both HS-induced CD25-CCR7 complex formation and integrin activation. Thus, both anti-CD25 antibody and HS require common features of IL-2 needed for CD25-CCR7 complex assembly and resulting integrin activation. Collectively, these data show that IL-2 promotes CD25 interaction with CCR7, thereby forming the signal initiating complex. Furthermore, canonical and alternative IL-2 signaling can be decoupled by an IL-2 mutation, creating a tool to specify the biological role of alternative IL-2 signaling in immune responses.

## Introduction

Interleukin-2 (IL-2) controls the expansion and maintenance of T cell subsets. IL-2 binds to the α receptor (CD25), encoded by *IL2RA*, to promote IL-2 association with the signaling β(CD122) and γ(CD132) subunits; the resulting trimeric receptor triggers canonical signaling via associated Janus Kinases (JAK) 1 and 3 that phosphorylate Signal Transducer and Activator of Transcription (STAT)5 (1, 2). IL-2 also signals through a heterodimeric complex comprising the β and γ subunits, albeit with a 100-fold lower affinity (3). CD4^+^FOXP3^+^CD25^Hi^ regulatory T cells (Tregs), which provide a critical brake on immune responses and inflammation(4), are dependent on IL-2 and integrins. Dynamic increases in integrin affinity for macromolecular ligands (5)(“activation”) are essential for the function and phenotype of Tregs (6). Thus, augmenting integrin activation in Tregs may enhance suppressive function and blunt autoimmunity and inflammation.

PC61 (7), in contrast with other anti-mouse CD25 antibodies, induces integrin activation in Tregs and amplifies their suppressive function (8). PC61 binding to CD25, which lacks endogenous signaling capability, induces a complex formed by the IL-2 dependent association of CD25 with CCR7. In contrast to signaling via STAT5, PC61 induces Gαi/o-dependent integrin activation. Heparan sulfate (HS) docks IL-2 in tissues (9) and can promote Treg suppressive function(10, 11). HS biases IL-2 signaling towards this Gαi-dependent pathway(8). We termed this alternative pathway **M**onoclonal Antibody (or **M**atrix)-Directed **A**lternative **C**ytokine **S**ignaling (MACS)(8).

Here, we characterize the mechanism whereby CD25 associates with CCR7 to trigger MACS. Structure-based design coupled with combinatorial screening identified an IL-2 mutant (IL-2(E52K)) that leaves canonical IL-2 signaling intact but disrupts MACS-induced integrin activation. The identification of such a mutant indicates that IL-2 facilitates association of CD25 to CCR7, thereby initiating MACS. We further report that heparin sulfate (HS), a physiological activator of MACS (8), induced IL-2 dependent CD25-CCR7 association and consequent integrin signaling that was lost in IL-2(E52K). In sum, this work establishes the initiation mechanism for the IL-2-driven MACS pathway in humans, shows that anti-CD25 antibodies and HS induce MACS through common features of IL-2, and provides a new tool to analyze the biological consequences of MACS.

## Results

### IL-2 promotes interaction of CD25 with CCR7

IL-2 is an obligatory component of the PC61-induced complex of CD25 and CCR7 (8). We reasoned, by analogy to IL-2-mediated assembly of the trimeric receptor (12), that PC61 causes CD25-bound IL-2 to interact with CCR7 to initiate MACS. We used the structure of trimeric receptor-bound IL-2 (12) to create IL-2 mutants that do not disrupt interaction with the αβγ receptor. We selected charge reversal substitutions in unstructured loops to minimize potential protein misfolding (Fig. 1A). With these considerations, we constructed 10 such human IL-2 mutations (Table 1).

**Table 1.** Mutations in human IL-2.

| Mutant # | Mutated residue | Mutation site with flanking sequence |
| --- | --- | --- |
| 1 | Q22V, Q126A, I129D, S130G | LDLVMIL,<br>TFCASID**G**TLT |
| 2 | Y31A, K32E | INNA**E**NPK |
| 3 | K76E, F78A, H79A | AQSENAALRP |
| 4 | K8D, K9D | SST**DD**TQL |
| 5 | K48D, K49D | YMP**DD**ATE |
| 6 | E52K | KATKLKH |
| 7 | D109K, E110K | EYAK**K**TAT |
| 8 | R81E | FHLEPRD |
| 9 | D84K | RPR**K**LIS |
| 10 | R83E | LRPEDLI |

IL-2 mutants and the sites of mutated amino acid residues. Mutations are in bold text.

**Figure 1.**
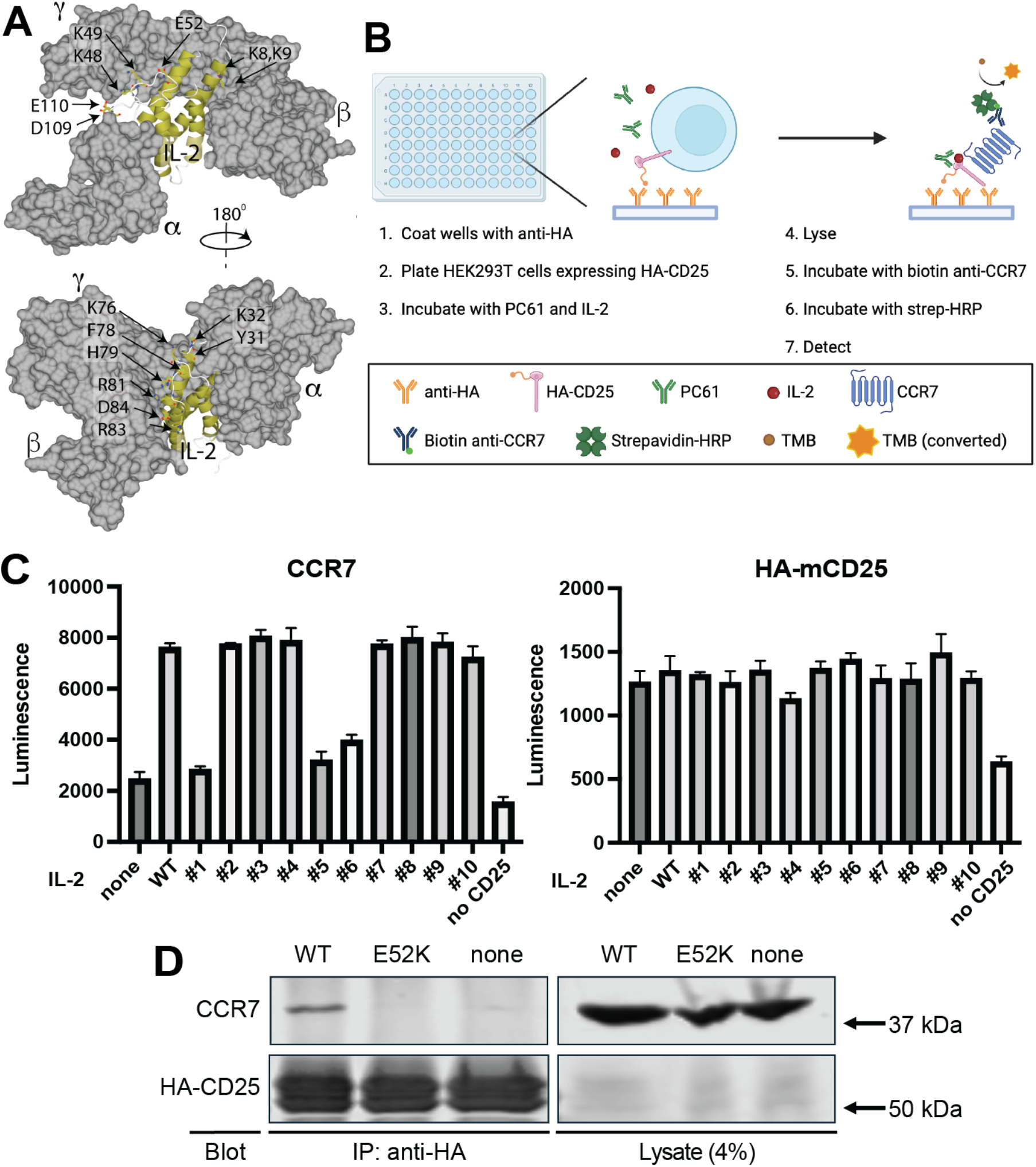
IL-2(E52K). does not support CD25-CCR7 complex. (A) *Design Strategy*. Depicted are space filling models of the trimeric IL-2 receptor (gray) with the helical elements of IL-2 depicted as a gold ribbon. Charged residues in unburied IL-2 loops are labelled and are targeted for mutagenesis. (B) *Schematic of ELISA*. HEK293T-HA-mCD25 cells were plated on anti-HA-coated wells and then incubated with 1 μg/mL IL-2 (red) and 5 μg/mL PC61 (green) for 1 h at 37 ^o^C. Cells were lysed and washed plates were probed by anti-CCR7 or anti-HA followed by chemiluminescence reagents. (C) *ELISA screen identifies mutants that fail to support CD25-CCR7 complex*. Note that CCR7 association is IL-2 dependent and reduced in mutants 1,5, and 6. (Mean ± SEM. N=3). D) *IL-2(E52K) fails to support CD25-CCR7 complex*. Experiment was performed as in panels B and C with detection by immunoprecipitation of HA-CD25 followed by immunoblotting.

An enzyme-linked immunosorbent assay measured the association of CD25 to CCR7 with detection by chemiluminescence (Fig. 1B). HEK293T cells expressing HA-tagged mouse CD25 (mCD25), were plated on anti-HA coated plates and were incubated with PC61 and IL-2, followed by cell lysis and measurement of the bound CCR7 and CD25. The increase in luminescence when both IL-2 and PC61 were added to mCD25-expressing cells revealed that PC61 induced IL-2-dependent CD25-CCR7 association, whereas this increased luminescence was not observed with addition of PC61 or IL-2 alone. IL-2 mutants 1, 5, and 6 showed a two-fold reduction in luminescence; we selected 6 (IL-2(E52K)) for further study because mutant 1 is known to block γ subunit association (13) and 5 was not well-secreted. Co-immunoprecipitation of CCR7 and mCD25 confirmed that IL-2, but not IL-2(E52K), enables PC61-induced CD25-CCR7 association (Fig. 1D).

We used HEK293T cells expressing human CD25 (hCD25), to screen for anti-hCD25 antibodies that drive CD25-CCR7 association. Among these, we found that 7G7B6, but not BC96, induced IL-2-dependent hCD25-CCR7 association (Fig. 2A); in sharp contrast, IL-2(E52K) failed to support 7G7B6 induced hCD25-CCR7 complex (Fig. 2B). Importantly, IL-2(E52K) did not impair IL-2 binding to CD25(Fig. 2C). Thus, specific features of IL-2 promote association of CD25 to CCR7 in the presence of certain anti-CD25 antibodies.

**Figure 2.**
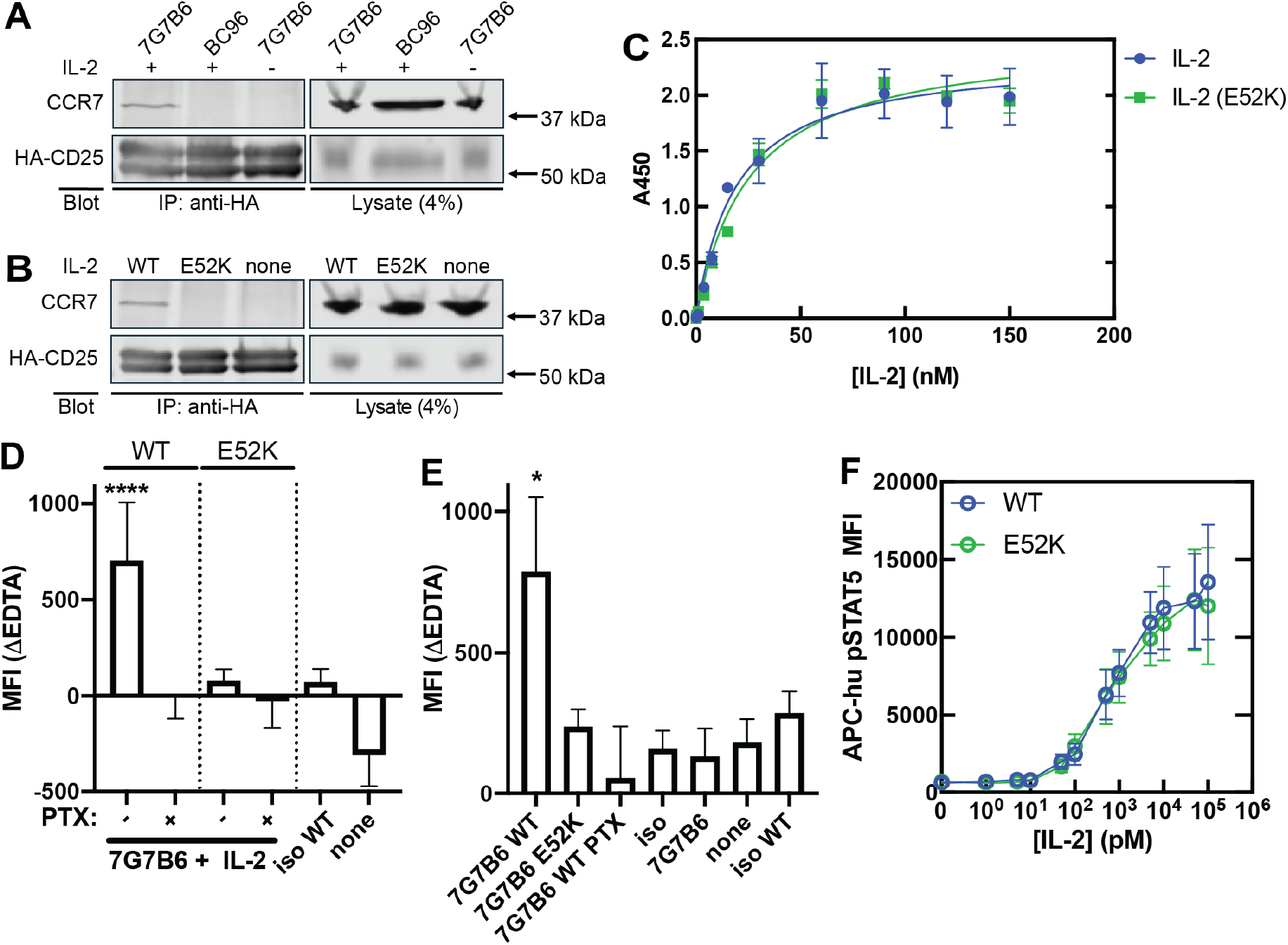
Assembly of human CD25-CCR7 complex triggers integrin activation. (A) *7G7B6 triggers IL-2 dependent hCD25-CCR7 complex*. 5 ug/mL anti human CD25 antibodies 7G7B6 or BC96 were incubated with HA-hCD25 expressing HEK293T cells in the presence or absence of 1 ug/mL IL-2 and following lysis and immunoprecipitation with anti-HA were assayed for captured CCR7 or HA-hCD25 by immunoblotting. (B) *IL-2(E52K) does not support 7G7B6-induced hCD25-CCR7 complex*. IL-2(E52K) or wild type IL-2 were assayed for capacity to support 7G7B6-induced CD25-CCR7 complex as described in panel A. (C) *IL2(E52K) mutation does not disrupt CD25-IL-2 interaction*. Microplates coated with recombinant hCD25 extracellular domain were incubated with the indicated concentrations of IL-2 or IL-2(E52K) and bound IL-2 was quantified by ELISA. (D) *IL-2(E52K) does not trigger MACS-dependent integrin activation*. IL-2Rα YT-1 cells. were incubated with indicated reagents in the presence of 5 μg/mL hMAdCAM-1 Fc for 30 min at 37 ^o^C, followed by APC-anti-human IgG Fc and FACS to assess MAdCAM-1 binding. (E) *IL-2(E52K) does not trigger MACS-dependent integrin activation* Freshly isolated human Tregs were incubated with 7G7B6 and 5 μg/ml VCAM-1 Fc (these cells express little α4β7) and 7G7B6 in the presence or absence of IL-2(WT) or IL-2(E52K) (E52K) for 30 min at 37 ^o^C before measurement of bound VCAM-1 Fc by FACS. In some experiments 200 ng/ml Pertussis toxin was added (PTX) or an irrelevant antibody (iso) was added in place of 7G7B6. *(F) IL-2(E52K) triggers canonical IL-2 signaling*. IL2Rα YT-1 cells were stimulated with varying concentrations of IL-2 or IL-2(E52K) for 30 min at 37 ^o^C, followed by staining with APC-anti phospho-STAT5 and analysis by FACS. *: p<0.05, **: p<0.01, NS: not significant by one-way ANOVA.

### IL-2(E52K) blocks MACS-dependent integrin activation

In the presence of 7G7B6, IL-2 stimulated increased MAdCAM-1 (Fig. 2D) binding, a measure of activation of integrin α4β7(14), to an immortalized CD25^Hi^ NK cell line, IL-2Rα^+^ YT-1 (15, 16). Increased MAdCAM-1 binding was blocked by Pertussis toxin, confirming it was Gαi-dependent. In sharp contrast, IL-2(E52K) supported markedly reduced MAdCAM-1 binding. These results were recapitulated with primary human CD25^Hi^CD127^Lo^ Tregs, wherein we observed that in the presence of 7G7B6, administration of IL-2 but not IL-2(E52K) increased integrin activation (Fig. 2E). Consistent with preservation of CD25 binding in IL-2(E52K), IL-2 and IL-2(E52K) induced phosphorylated STAT5 (pSTAT5) at comparable concentrations (Fig. 2F), indicating that the E52K mutation did not inhibit the canonical JAK/STAT-dependent IL-2 signaling pathway. Thus, IL-2 MACS can be selectively disabled by the IL-2(E52K) mutation.

### Heparan sulfate (HS) triggers IL-2-dependent interaction of CD25 and CCR7, resulting in MACS

We previously showed that heparan sulfate (HS), a physiological ligand of IL-2, can bias IL-2 signaling towards integrin activation (8). Incubation of solubilized membranes of hCD25-expressing HEK293T cells with HS and IL-2, but not IL-2(E52K), resulted in co-immunoprecipitation of CD25 and CCR7 (Fig. 3A-B). Furthermore, in the presence of IL-2, addition of HS to IL-2Rα^+^ YT-1 cells, caused increased MAdCAM-1 binding that was blocked by Pertussis toxin (Fig. 3C). In contrast, IL-2(E52K) failed to support HS-induced MAdCAM-1 binding. At this concentration, HS did not alter the pSTAT5 response to IL-2 (Fig. 3D). These data show that HS, like certain anti-CD25 antibodies, can trigger IL-2-dependent CD25-CCR7 association that results in MACS.

**Figure 3.**
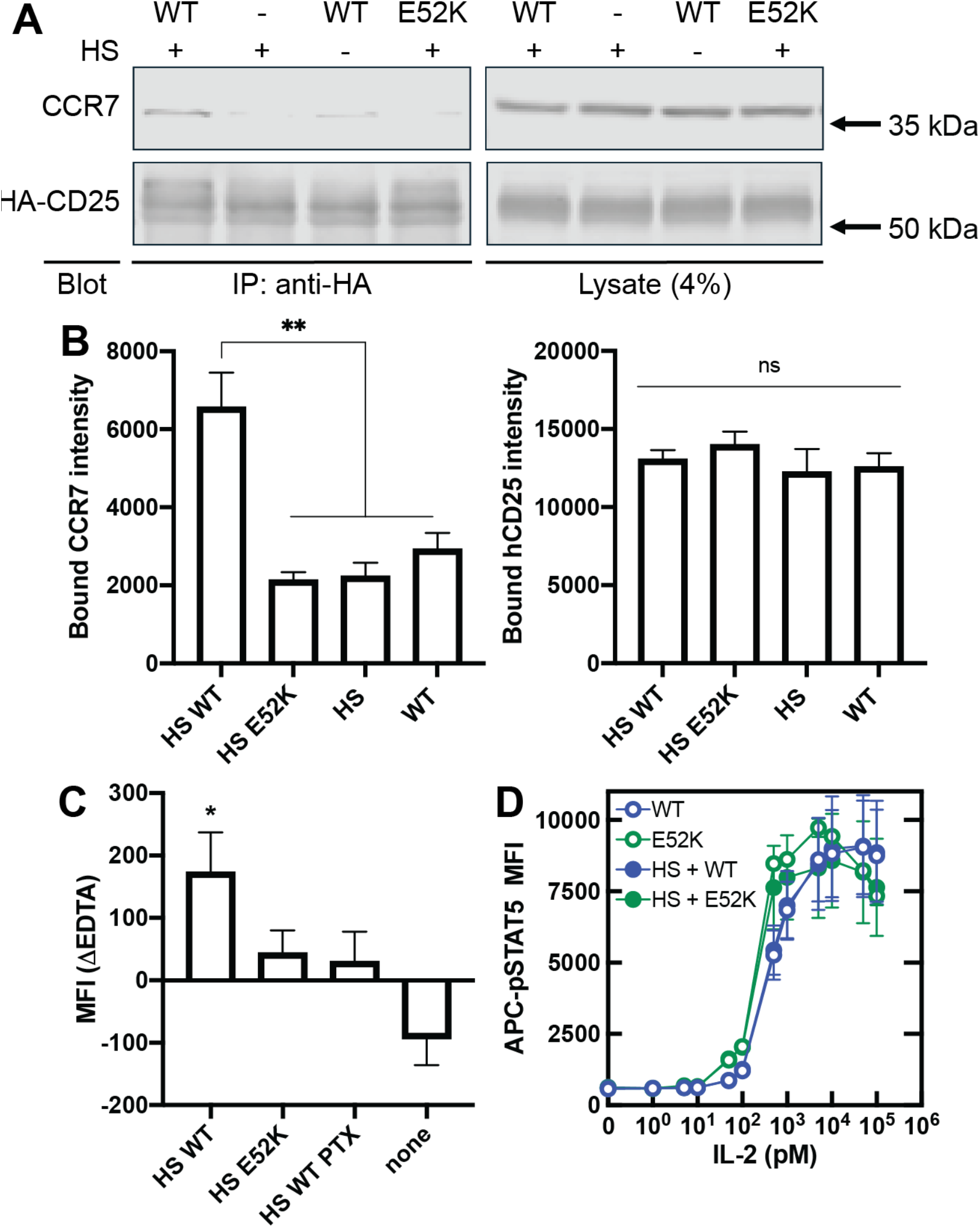
HS induces an IL-2 dependent CD25-CCR7 complex to trigger MACS. (A) *IL-2(E52K) does not support HS induced CD25-CCR7 association*. Membranes from HA-CD25 expressing HEK293T cells were incubated with 5 μg/mL HS in the presence or absence (-) of 2.5 μg/mL IL2 (WT) or IL-2 (E52K (E52K) for 30 min at 37 ^o^C. Anti-HA immunoprecipitates were fractionated by SDS-PAGE and blots were probed with either anti-HA or anti CCR7. (B) *Quantification of data from experiments performed as in panel B*. Blots from 3 such experiments were quantified by Licor infrared spectroscopy. (C) *IL-2(E52K) does not support HS induced integrin activation*. IL2Rα YT-1 cells were incubated with 5 μg/mL HS in the presence or absence (-) of 0.5 μg/mL IL2 (WT) or IL-2(E52K) (E52K) and 5 μg/ml MAdCAM-1 for 30 min at 37°C and bound MAdCAM-1 was measured by FACS. None signifies no additions. (D) *5 μg/mL HS did not affect STAT5 phosphorylation*. IL2Rα YT-1 cells were incubated with varying concentrations of IL-2 or IL-2(E52K) in the presence or absence of 5 μg/ml HS for 30 min at 37° followed analysis of phsopho-STAT5 staining by FACS. *: p<0.05, **: p<0.01 by one-way ANOVA.

## Discussion

IL-2 signaling via the αβγ receptor shapes immune responses by promoting the proliferation of T cells and, in particular, enabling the development, function, and stability of Tregs (1). In the presence of an anti-mCD25 antibody, PC61, IL-2 can induce activation of Treg integrins associated with formation of a mCD25-CCR7 complex. Here, we used the structure of IL-2 bound to the canonical receptor to construct an IL-2 mutant (IL-2(E52K)) that fails to support the PC61-induced mCD25-CCR7 complex. We identified an anti-hCD25 antibody, 7G7B6, that induces a hCD25-CCR7 complex in the presence of IL-2 but not IL-2(E52K). Notably, IL-2(E52K) failed to support 7G7B6-induced integrin activation both in human CD25-expressing cells and in human primary Tregs. Nevertheless, IL-2(E52K) and wild type IL-2 bound to CD25 with similar affinity and induced signaling via the canonical αβγ receptor with similar potency. Collectively, these data indicate that IL-2 promotes CD25 interaction with CCR7, thereby initiating MACS. Moreover, our IL-2 mutant serves as a molecular probe that can decouple canonical and MACS IL-2 signaling. HS, which is a physiological IL-2 ligand (9), promote IL-2-induced integrin activation (8). Here, we found that HS-induced IL-2 dependent hCD25-CCR7 association was lost with IL-2(E52K), corresponding with reduced integrin activation. In addition to illuminating the mechanism of IL-2 signaling via CCR7, these data establish that HS can trigger a CD25-CCR7 complex that, analogous to certain anti-CD25s, which triggers MACS.

The results of this work indicate that IL-2 MACS results from the interaction of IL-2-bound CD25 with CCR7. The anti-mouse CD25 antibody PC61 triggers CD25-CCR7 association and integrin activation; and both responses are inhibited by an anti-IL-2 antibody that blocks binding to CD25(8). Furthermore, a CD25 mutant that disrupts the IL-2 binding site prevents assembly of the anti-CD25-induced CD25-IL-2-CCR7 complex (8); thus complex formation requires IL-2-CD25 interaction. Here we find that an IL-2 mutation that spares the CD25 binding site but disrupts the CD25-IL-2-CCR7 complex. Furthermore, our study demonstrates that formation of the CD25-CCR7 complex triggers MACS, as an IL-2 mutation that disrupts the complex also suppresses integrin activation. Importantly, this mutation did not affect IL-2 binding to CD25 or IL-2-induced STAT5 phosphorylation, showing that MACS can be disabled while canonical IL-2 signaling is maintained. The fact that IL-2(E52K) disrupts MACS but not canonical IL-2 signaling suggests that IL-2 may serve as a bridge between CD25 and CCR7, possibly directly interacting with both receptors in a ternary complex that may or may not involve receptor/receptor contact. However, we acknowledge the possibility that IL-2 may allosterically modulate CD25 to enhance interaction with CCR7, and the interaction may be exclusively between the receptors. Structural studies will elucidate the molecular details of complex formation.

HS is a physiological ligand for IL-2 and can enhance IL-2-induced integrin activation (8). We report that HS-induced integrin activation is associated with formation of a CD25-CCR7 complex and disruption of this complex by the IL-2(E52K) mutation blocks integrin activation. Thus, HS-triggered MACS can be ascribed to HS enabling IL-2 to mediate assembly between CD25 and CCR7. These results will lead to new work to discern how HS and certain anti-CD25 antibodies can enable IL-2 to connect CCR7 and CD25 to trigger MACS. Furthermore, IL-2 mutants that selectively disrupt MACS and spare canonical IL-2 signaling, such as IL-2(E52K), will enable studies to define the role of this newly described signaling pathway in immune homeostasis and immunopathogenesis.

## Experimental Procedures

### Reagents and Antibodies

Anti-CCR7 polyclonal antibody (Novus Biologicals #NBP2-67324), anti-HA mouse monoclonal antibody (12CA5), anti-HA mouse monoclonal antibody (BioLegend #900801), anti-HA rabbit polyclonal antibody (BioLegend #923501), biotin-anti-human CCR7 mouse monoclonal antibody (BioLegend #353240), biotin-anti-human CD25 goat polyclonal antibody (R&D Systems # BAF223), biotin-anti-mouse CD25 goat polyclonal antibody (R&D Systems # BAF2438), anti-mouse CD25 monoclonal antibody (clone PC-61.5.3, BioXcell # BE0012), anti-human CD25 monoclonal antibody (7G7B6)(17), anti-human CD25 monoclonal antibody (BC96, BioLegend #302609), Streptavidin-horseradish peroxidase (Invitrogen, #S911), IRDye® 800CW Goat anti-Mouse IgG Secondary Antibody (LI-COR 926-32210), IRDye® 800CW Goat anti-Rabbit IgG Secondary Antibody (LI-COR #926-32211), IRDye® 680RD Goat anti-Rabbit IgG Secondary Antibody (LI-COR #926-68071), IRDye® 680RD Goat anti-Mouse IgG Secondary Antibody (LI-COR #926-68070), Protein G agarose (GenScript # L00209).

### Cell Culture

HEK293T cell lines expressing HA-tagged mouse CD25 or human CD25 were constructed and cultured as described (14). IL-2 Rα^+^ YT-1 cells were derived from YT-1 cells (16), induced to express endogenous CD25, as described previously (15).

### ELISA

The formation of the CD25-CCR7 complex was assayed by ELISA as previously described (8). Briefly, Microplate wells were coated with anti-HA antibody and HEK293T cells expressing HA-CD25 were added to the plate and stimulated with 5 μg/ml PC61 and 1 μg/ml IL-2 for 1h at 37ºC. Cells were lysed and the plate was then washed and bound CCR7 or CD25 were labeled with biotinylated primary antibodies and after washing bound antibodies were complexed with streptavidin-horse radish peroxidase followed by chemiluminescence substrate (Thermo Scientific, #32106). Chemiluminescence intensity was assessed by microplate scanner (Spark®, Tecan Life Sciences) to measure bound CCR7. In subsequent experiments bound anti-CD25 or CCR7 were assessed by SDS-PAGE followed by immunoblotting.

### Measurement of IL-2 binding to CD25

The 6x-His tagged extracellular domain of human CD25 (amino acid residues 1-217), human IL-2, and the IL-2(E52K) were expressed in Expi293 cells (Thermo Fisher) and purified using His•Bind® Resin (EMD Millipore). Microplates (Greiner Bio-One) were coated with hCD25 (2μg/ml) in carbonate/bicarbonate coating buffer (pH 9.2) and blocked with 2% BSA. Increasing concentrations of IL2 or IL-2(E52K) were added, incubated for 2 hours at RT, and then washed. Subsequently, the bound IL-2 was quantified using the ELISA MAX™ Deluxe Set Human IL-2 (BioLegend).

### Subcellular Fractionation, Co-Immunoprecipitation, and Immunoblot Analysis

Membrane fractions from HA-CD25 expressing HEK293T cells were prepared by differential centrifugation as described (18). Cell membranes that were washed and then suspended in fractionation buffer containing 1% Nonidet P-40 on ice for 1 hr. The solubilized membrane preparations were incubated with 5 μg/mL heparan sulfate at 37 ^o^C for 30 minutes and incubated with anti-HA antibody immobilized on protein G beads in the lysis buffer at 4 ^o^C for 2 hrs to overnight to capture HA-CD25. The isolated protein complex was washed with the lysis buffer and subjected to SDS-PAGE. Bound proteins were detected by immunoblotting with designated antibodies.

### Isolation and Activation of Human Primary CD4^+^ T Lymphocytes

Human blood was obtained from normal adult human volunteers. Red blood cells from the acquired whole blood were lysed with ACK lysis buffer (Gibco #A1049201), followed by magnet separation using human whole blood CD4+ T cells negative isolation kit (BioLegend #480162). For soluble ligand binding assays with freshly isolated CD4+ T lymphocytes, cells were labeled with PerCP-hCD4 (BioLegend #357414), FITC-hCD127 (Biolegend #351312), and PE-hCD25 (BioLegend #302606) to distinguish CD25^hi^CD127^lo^ Tregs and Tconv.

### Soluble Ligand Binding Assay

Cells were plated in a 96-well plate and incubated with the following ligands for 30 min at 37 ^o^C in HBSS with Ca/Mg (Gibco #14-025-092): 5 μg/mL human MAdCAM-1 Fc (R&D Systems #6056-MC) for YT-1 hCD25 cells or human VCAM-1 Fc (R&D Systems #862-VC) with human primary CD4^+^ T cells, 0.5 μg/mL IL-2, 10 μg/mL IgG control or 7G7B6, 5 μg/mL Heparan Sulfate, 200 ng/mL Pertussis toxin, 100 nM PMA (Sigma #P8139), and 10 mM EDTA (Invitrogen #AM9260G). Cells were then immediately washed with 1X HBSS with Ca/Mg, followed by incubating with APC-anti-human IgG Fc (1:25, Biolegend #410714) for 1 h on ice. Cells were washed in 1X HBSS with Ca/Mg, and geometric mean fluorescence intensity (MFI) was determined on an Accuri C6+ flow cytometer (BD Biosciences). Geometric mean fluorescence intensity (MFI) of EDTA groups were subtracted from MFI without EDTA groups to report ligand binding (MFI ΔEDTA). Triplicate measurements were made in each experiment and each experiment was independently replicated at least three times with similar results.

### STAT5 Phosphorylation

Cells were plated in each well of a 96-well plate and incubated with indicated concentrations of IL-2, or 5 ug/mL HS in 1X PBS. Cells were stimulated for 30 min at 37°C and immediately fixed and permeabilized using the Transcription Factor Phospho Buffer Set (BD Biosciences #563239). Cells were then washed and incubated with APC-anti human STAT5 (1:50, BD Biosciences #612599) for 1 h on ice in the wash buffer. Cells were then washed twice, once with the wash buffer and next with 1X PBS, and MFI was determined on Accuri C6+ flow cytometer (BD Biosciences). Duplicate measurements were made in each experiment and each experiment was independently replicated at least three times with similar results.

### Statistical Analysis

Statistical analysis was performed using PRISM software (version 9.00, GraphPad Software), and all datasets were checked for Gaussian normality distribution. Data analysis was performed using two-tailed t-test, one-way ANOVA or two-way ANOVA followed by Bonferroni post-test as indicated in the Fig. legends. The resulting P values are indicated as follows: NS: not significant, *: p <0.05; **: 0.01< p <0.05, ***: 0.001< p <0.01, ****: p <0.001. Plotted data are the mean ± SEM of at least three independent experiments.

All data are contained within the manuscript.

Supported by P01 HL 15433 (MHG, KL). The content is solely the responsibility of the authors and does not represent the official views of the National Institutes of Health.

The authors declare that they have no conflicts of interest with the contents of this article.

## Notes

### Competing Interest Statement

The authors have declared no competing interest.

